# Genuine Directed Evolution In Test Tube (GENie)

**DOI:** 10.64898/2026.05.04.722721

**Authors:** Lilin Feng, Maochao Mao, Ulrich Schwaneberg

## Abstract

Directed evolution has long been constrained by complex screening hardware and labor-intensive workflows. Here, we report the first genuine *test-tube* screening platform that uses His6-tagged peptide–functionalized magnetic beads and Fe^3+^-decorated *E. coli* cells to establish a phenotype–genotype linkage, thereby decoupling ultrahigh-throughput screening from specialized instrumentation and democratizing directed evolution. The platform demonstrated a screening throughput of > 10^8^ events s^−1^ and an enrichment factor of up to 63-fold. Using galactose oxidase as a model, we identified variants with up to a 26-fold increase in catalytic efficiency. Extensions to D-amino acid oxidase and alcohol oxidase yielded variants with up to 5383-fold and 25-fold improvements over their respective wildtypes after a single round of screening. These results highlight the platform’s capacity to rapidly engineer H_2_O_2_-generating oxidases and to advance AI-driven enzyme design through rapid data generation.

## INTRODUCTION

Directed evolution allows scientists to engineer enzymes and binding proteins through iterative cycles of mutation and selection, bypassing the need for a detailed mechanistic understanding of sequence–function relationships(*1, 2*). A fundamental challenge in directed evolution lies in the fact that nature provides 20 different building blocks for each position in a protein. Consequently, even a relatively small protein of 100 amino acids corresponds to a theoretical sequence space of 20^100^ (>10^130^) variants. Identifying rare improved variants within this immense sequence space requires massive screening efforts, which have historically constrained directed evolution to reliance on high-throughput instrumentation and labor-intensive screening workflows(*3, 4*). Although conceptually ambitious, the realization of a genuine *test-tube* screening platform that minimizes dependence on complex instrumentation would substantially reduce costs, simplify workflows, and accelerate experimental cycles. Lowering these technical barriers would facilitate rapid hypothesis testing, enable more frequent iterative cycles, and support parallel exploration across diverse conditions, thereby broadening access to directed evolution and empowering its applications in biocatalysis, drug discovery, and synthetic biology across a wide range of laboratory settings.

Realizing this vision, however, requires the development of a genuine *test-tube* screening platform that meets four key criteria: (i) robust genotype–phenotype linkage to accurately associate enzyme function with its encoding sequence; (ii) sufficiently high throughput to effectively sample large libraries; (iii) high enrichment efficiency to isolate rare beneficial variants; and (iv) broad applicability across diverse enzyme classes(*5*). Currently, flow cytometry and chip-based microfluidic screening systems represent the two principal ultrahigh-throughput screening (uHTS) formats in directed enzyme evolution (*5–10*). Flow cytometry enables rapid analysis of millions of variants by correlating fluorescent signals with individual cells producing active variants, but relies on sophisticated instrumentation and suffers from moderate enrichment due to product diffusion-mediated crosstalk (*11–13*). In contrast, microfluidic systems minimize crosstalk through physical compartmentalization, achieving higher enrichment efficiency, albeit at the cost of increased technical complexity and specialized device fabrication (*7, 14*).

To decouple screening from complex instrumentation, magnetic beads provide a simple and effective platform for biomolecular compartmentalization and separation (*15–18*). Typically based on superparamagnetic iron oxide nanoparticles (SPIONs), these magnetic beads feature a high surface-to-volume ratio for efficient molecular binding and can be rapidly and reversibly manipulated by an external magnetic field(*19*). Their compatibility with standard and scalable laboratory workflows has led to widespread use in diagnostics and biomolecule purification(*20*). To harness the magnetic carriers for directed evolution, material-binding peptides (MBPs) provide a versatile, genetically encoded anchoring strategy (*21*). MBPs, typically 20-100 amino acids in length, exhibit strong affinity for diverse natural and synthetic material surfaces under mild conditions, driven by non-covalent multiple-site interactions (e.g., electrostatic, hydrophobic, π-π stacking, and hydrogen bonding). Notably, recent studies have demonstrated that the 47-amino-acid MBP LCI (liquid chromatography peak I) from *Bacillus subtilis* binds strongly to SPIONs, providing a foundation for the development of a genuine test-tube screening platform (*22, 23*).

Here, we introduce a genuine test-tube screening platform that leverages magnetic capture for ultrahigh-throughput screening of large mutant libraries (Fig. 1). In this approach, *E. coli* cells expressing an oxidase library are incubated with substrate and Fe^2+^ in a single pot. During the oxidation reaction, H_2_O_2_—a by-product of the oxidation reaction—diffuses across the cell envelope and drives local oxidation of Fe^2+^ to Fe^3+^ via the Fenton reaction (Fe^2+^ + H_2_O_2_ → Fe^3+^ + HO.+ HO^−^), the produced Fe^3+^ then deposits on the surface of *E*.*coli* cells expressing active oxidase mutants. These cells selectively assemble with Fe_3_O_4_ nanoparticles functionalized with His_6_-tagged LCI through specific coordination between the His_6_-tag and cell surface-deposited Fe^3+^, thereby establishing a robust genotype–phenotype linkage that enables rapid magnetic separation. The Genie platform demonstrated a screening throughput of > 10^8^ events s^−1^ and an enrichment factor of up to 63-fold. Using galactose oxidase (GalOx) as a model, we validated the platform under two evolutionary pressures targeting substrate affinity and catalytic turnover, identifying two variants with up to 26-fold improvement over wildtype, among which G2 (*K*_m_) showed a 5.8-fold decrease in *K*_m_ and G2 (*k*_cat_) showed a 7.8-fold increase in *k*_cat_ than wildtype. The platform was further extended to D-amino acid oxidase (D-AAO) and alcohol oxidase (AOx), identifying variants with improvements of up to 5383-fold and 25-fold over their respective wildtypes after a single round of screening, demonstrating its general applicability and ability to efficiently isolate rare beneficial variants without specialized instrumentation.

**Fig. 1.**
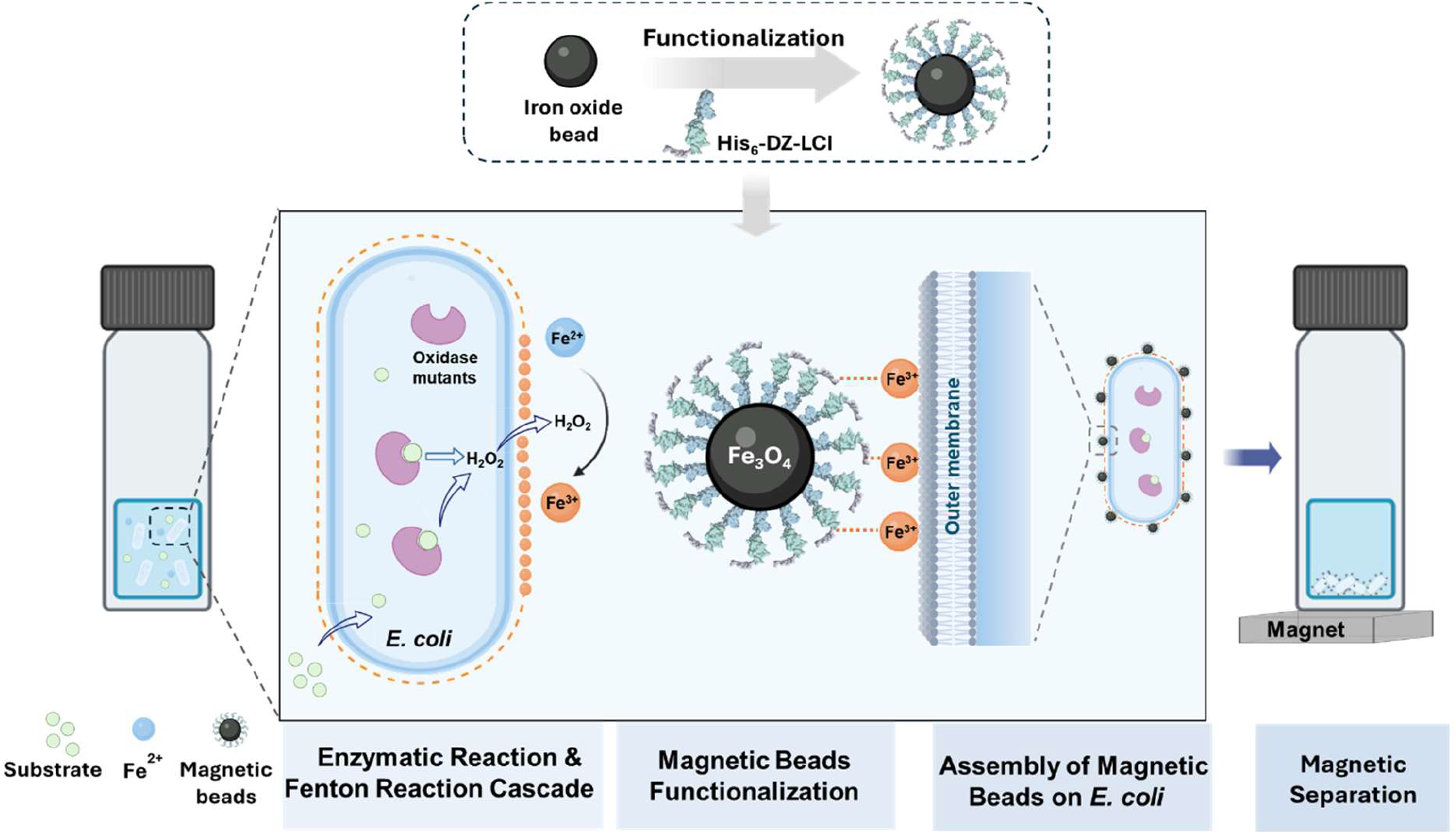
Schematic representation of the GENie screening system. First, *E. coli cells* expressing an oxidase library are incubated with substrate and Fe^2+^ in one pot. The generated H2O_2_ diffuses outside the cell envelope and oxidizes Fe^2+^ to Fe^3+^, which can be deposited on the cell surface. Subsequently, His_6_-LCI functionalized Fe_3_O_4_ beads are added and assembled on the *E. coli* cell surface, coordinated by the His_6_ tag and Fe^3+^ interaction. Afterward, magnetic separation can be performed to identify improved oxidase variants.

## RESULTS

### SPIONs functionalization

Previous studies have demonstrated that Fe^3+^ can deposit on *E. coli* cell surface and mediate the surface immobilization of His_6_-tagged proteins (*24–26*). Based on this observation, we hypothesized that cell surface-deposited Fe^3+^ could serve as an intermediary for the robust assembly of His_6_-tag-functionalized magnetic beads onto the *E. coli* cell surface. To validate the concept, we first functionalized Fe_3_O_4_ beads with the protein His_6_-DZ-LCI. A schematic illustration of the His_6_-LCI@Fe_3_O_4_ construct is shown in Fig. 2a. In this design, the iron oxide-binding peptide LCI anchors the construct to the Fe_3_O_4_ surface, while Domain Z (DZ) serves as a spacer that spatially separates the binding domain and exposes the His_6_-tag in the outer coating layer. Thermogravimetric analysis (TGA) indicates a mass loss increase from 5.6 % for bare Fe_3_O_4_ to 17.6 % after peptide functionalization, indicating successful attachment of His_6_-DZ-LCI. The derivative thermogravimetric (DTG) curve shows a characteristic peak at ~750 °C, further supporting the presence of peptide components on the SPION surface (Fig. 2b). Surface modification was also confirmed by X-ray photoelectron spectroscopy (XPS). The survey spectrum of His_6_-LCI@Fe_3_O_4_ shows the emergence of an N 1s signal together with an enhanced C 1s peak, both consistent with peptide coating (Fig. 2c). High-resolution N 1s spectra display two contributions at ~398.6 eV and ~400.3 eV, assigned to terminal –NH_2_ groups and amide (–CONH–) functionalities, respectively, further confirming peptide immobilization on the SPION surface (Fig. S1). Fourier-transform infrared spectroscopy (FT-IR) provides additional evidence for the modification. His_6_-LCI@Fe_3_O_4_ exhibits characteristic amide III, II, and I bands at 1260, 1540, and 1645 cm^−1^, respectively, along with weak C–H stretching signals at 2915 and 2984 cm^−1^, originating from aliphatic side chains of the peptide (Fig. 2d). The assembly of His_6_-LCI@Fe_3_O_4_ onto Fe^3+^-deposited *E. coli* cells was subsequently evaluated. We mixed the Fe^3+^-deposited *E. coli* suspension (OD_600_ = 2) with His_6_-LCI@Fe_3_O_4_ and performed magnetic separation; the *E. coli* cell suspension became immediately clear (Fig. 2e and Video S). In this regard, we calculated the cell capture efficiency with the throughput > 10^8^ events s^−1^. In contrast, *E. coli* suspensions without Fe^3+^ deposition or Fe_3_O_4_ without His_6_ tag did not exhibit magnetic separation under the same conditions (Fig. S2). Scanning electron microscopy (SEM) combined with energy-dispersive X-ray spectroscopy (EDXS) further confirmes the peptide coating on the SPION (indicated by red carbon signals) and the attachment of His_6_-LCI@Fe_3_O_4_ to the *E. coli* cell surface (indicated by Fe and O signals; Fig. 2f). Taken together, these results demonstrate the successful functionalization of Fe_3_O_4_ beads with His_6_-DZ-LCI and show that His_6_-LCI@Fe_3_O_4_ can be assembled on the *E. coli* cell surface through coordination between the His_6_-tag and Fe^3+^ ions.

**Figure 2.**
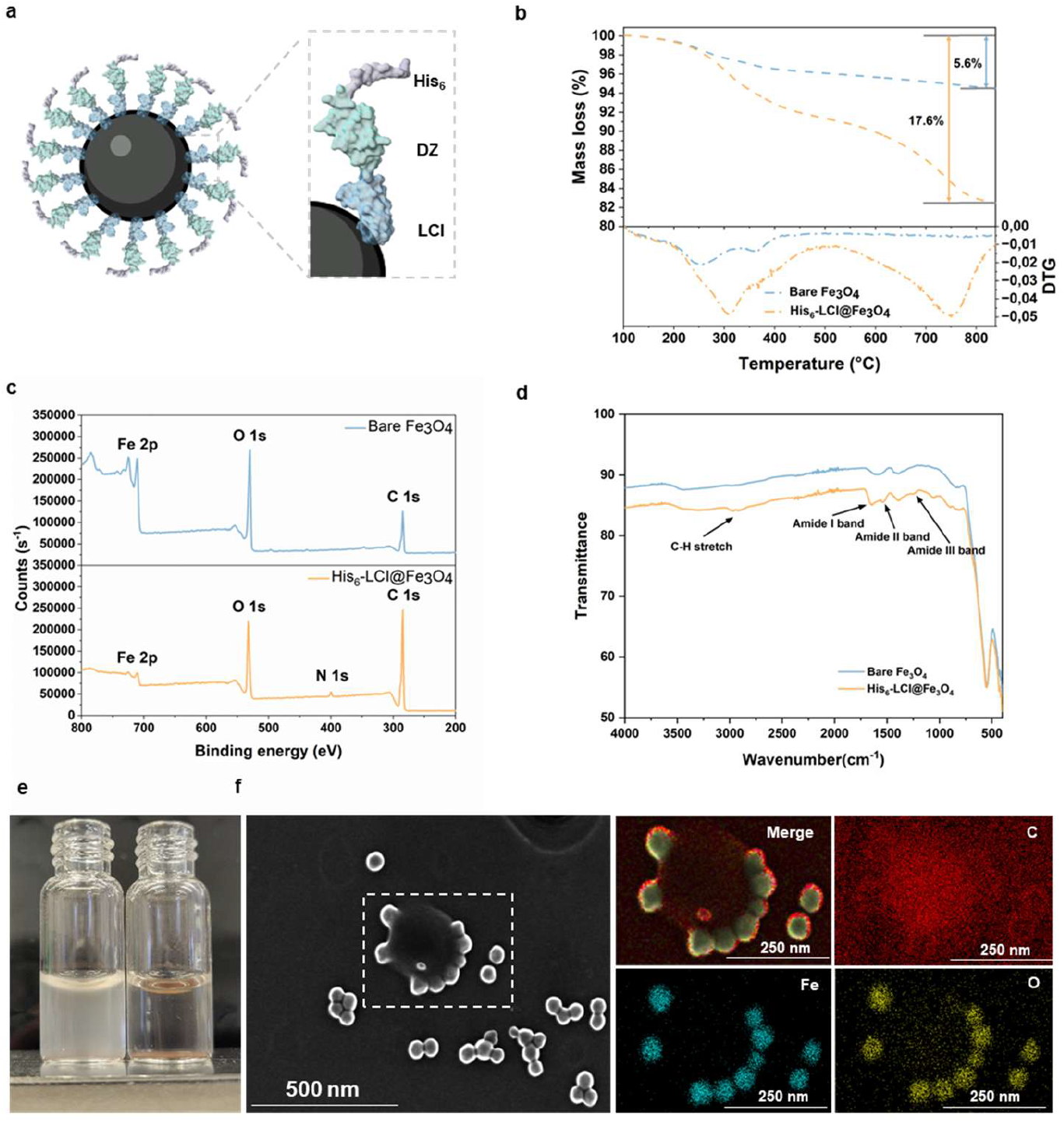
SPIONs functionalization with the peptide His_6_-DZ-LCI. a) Scheme illustrating decoration of a SPION by the peptide His_6_-DZ-LCI. b) TGA measurements of bare SPIONs (blue) and His_6_-LCI@Fe_3_O_4_ (orange). c) XPS spectra of bare SPIONs (top) and His_6_-LCI@Fe_3_O_4_ (bottom). d) FT-IR spectrum of bare SPIONs (blue) and His_6_-LCI@Fe_3_O_4_ (orange). e) Visualization of *E. coli* (OD = 2) before (left) and after magnetic extraction (right) by employing His_6_-LCI@Fe_3_O_4_ beads. f) SEM image of a single *E. coli* cell bound with His_6_-LCI@Fe_3_O_4_ beads, together with EDXS showing the merged signal and the distributions of C, Fe, and O, respectively. Protein models are visualized and coloured by ChimeraX 1.4.

### Enrichment assay developed by mCherry- and eGFP-expressing *E. coli*

Having demonstrated that *E. coli* cells displaying surface-bound Fe^3+^ can be captured by His_6_-LCI@Fe_3_O_4_ beads, we next evaluated the enrichment factor, defined as the fold increase in the fraction of target variants successfully recovered from a mixture after a single round of screening—a critical metric for the uHTS systems. To this end, we prepared the cell mixtures by combining mCherry-expressing *E. coli* with eGFP-expressing *E. coli* at ratios of 1:99, 5:95, and 10:90, among which the mCherry-expressing cells were pretreated with Fe^3+^. Fig. 3a shows the workflow developed for magnetic recovery of mCherry-expressing cells. Briefly, Fe_3_O_4_ nanoparticles were first functionalized by incubating them with the His-DZ-LCI peptide. After purification using magnetic separation, the resulting His_6_-LCI@Fe_3_O_4_ beads were incubated with the cell mixtures. The suspension was then shaken to promote attachment, followed by water bath sonication to eliminate nonspecific interactions. Subsequent magnetic separation was used to remove the supernatant and isolate bead-bound cells. The recovered cells were plated on auto-induction agar plates for qualitative visualization of enrichment, and flow cytometry was performed for quantitative analysis. Agar plate imaging demonstrated a distinct enrichment of rare red cell populations (mCherry-*E. coli*) from cell mixtures, indicated by a clear color shift from green (pre-enrichment) to red (post-enrichment) for both cell density conditions at OD_600_=0.1 and 0.01 (Fig. 3b). Flow cytometry analysis further revealed that, in the case of 1% mCherry-*E. coli*, the red cell population was enriched to approximately 63% after magnetic separation, corresponding to a ~63-fold enrichment. The enrichment became more pronounced for the 5% and 10% mCherry-*E. coli* cell mixtures, as evidenced by the enrichment of the red cell population to 89% and 92%, respectively (Fig 3c). These results demonstrate efficient recovery and enrichment of rare target cells from heterogeneous populations.

**Figure 3.**
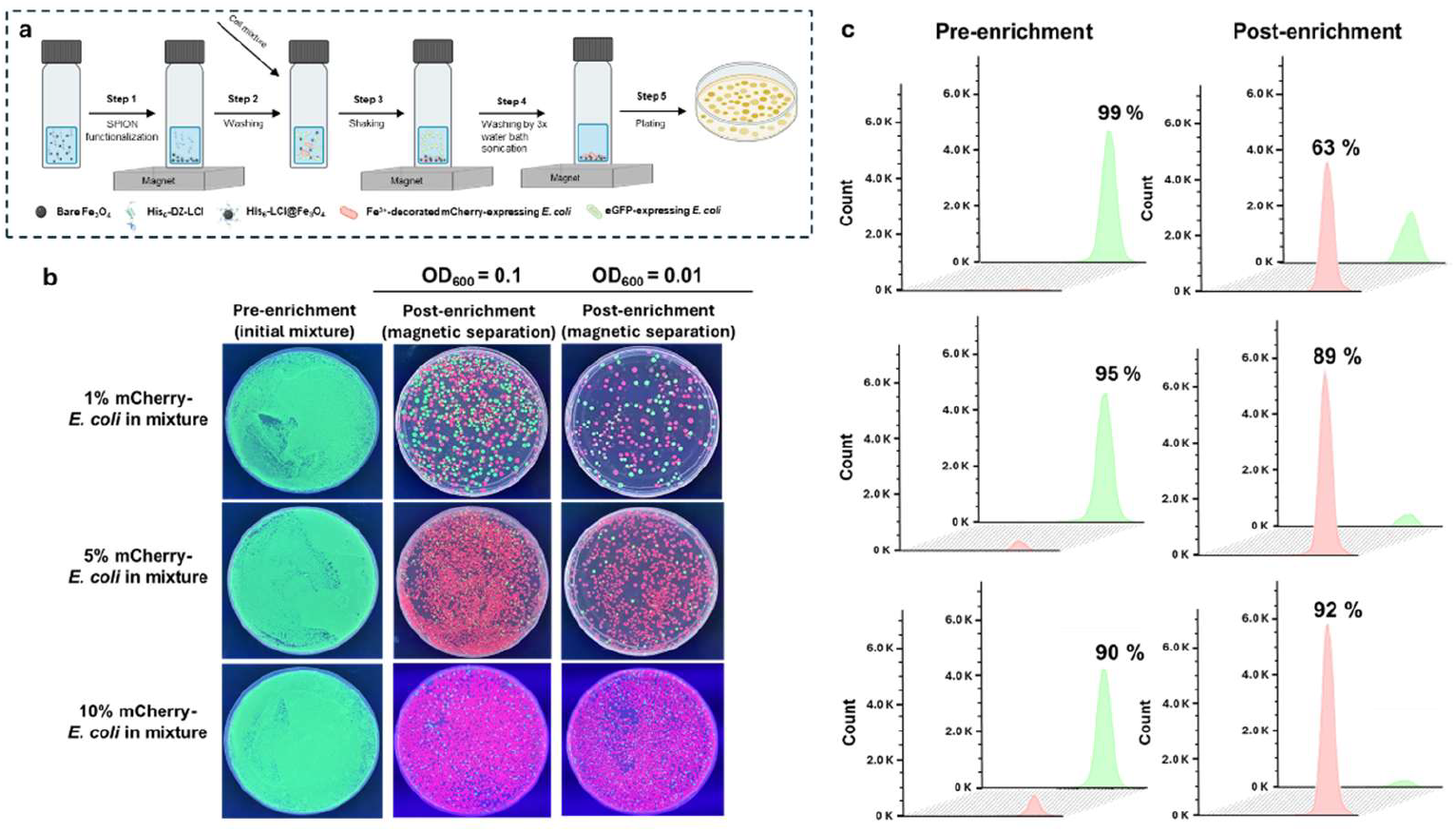
Proof of concept of the GENie platform for effective enrichment of target mCherry-*E. coli* cells from a cell mixture. (a) Cell mixtures are incubated with His_6__LCI@Fe_3_O_4_ beads, following the magnetic separation. (b) The auto-induction agar plates enable the quick visualization of the enrichment of target mCherry-*E. coli* cells before and after magnetic separation. In the right two columns, each plate displays the red and green colonies distribution after enrichment for two cell density conditions at OD_600_ = 0.1 (second column) and at OD_600_ = 0.01 (third column). The left column shows plates with initial mCherry-*E. coli* cells before enrichment. (c) Flow cytometry for the statistical evaluation of the mCherry-*E. coli* and eGFP-*E. coli* cells before and after magnetic separation (the total number of events is 150,000).

### Enrichment assay developed by GalOx- and eGFP-expressing *E. coli*

Next, we investigated whether the target *E. coli* cells expressing active oxidases could be selectively enriched from mixed populations using the GENie platform. We prepared the cell mixtures by combining the GalOx M1-expressing *E. coli* cells (GalOx M1-*E. coli*) with eGFP-expressing *E. coli* cells at ratios of 1:99, 5:95, and 10:90. We incubated the cell mixtures with galactose (the substrate of GalOx) and Fe^2+^ in one pot at ambient temperature. The reaction was terminated by the addition of 1,10-phenanthroline to chelate residual Fe^2+^ without affecting the coordination between Fe^3+^ and the His_6_-tag on the iron oxide bead surface, followed by magnetic separation to isolate GalOx M1-*E. coli* cells (Fig. 4a and S3). To minimize the background arising from Fe^2+^-mediated nonspecific binding of His_6_-LCI@magnetic beads, we systematically optimized the Fe^2+^ concentration by incubating *E. coli* cells (OD_600_ = 0.1) with varied Fe^2+^ concentrations in 0.9% NaCl (Fig. S4), and utilized His_6_-tagged eGFP to report the nonspecific binding. At high Fe^2+^ concentrations, we observed nonspecific labeling of His_6_-tagged eGFP on the cell surface, whereas at 60 µM Fe^2+^, no detectable fluorescence signal was present, indicating minimal background. The reaction time was further optimized to prevent overproduction of H_2_O_2_, which could diffuse and generate Fe^3+^ on neighboring inactive cells, leading to false positives. Following incubation, cell mixtures were centrifuged, and the morphology of the cell pellet was used as an indicator of sufficient Fe^3+^ deposition; cells bearing surface Fe^3+^ formed a compact, pointed pellet, in contrast to the rounded pellet of untreated cells (Fig. S5). This criterion was used to define appropriate reaction conditions for magnetic capture. The recovered cells were subsequently plated on auto-induction agar plates and analyzed by flow cytometry. Agar plate imaging exhibited a trend consistent with the aforementioned enrichment observed for red mCherry-*E. coli* cells. The significant enrichment of the rare GalOx M1-*E. coli* cells was evidenced by a clear color shift from green (pre-enrichment) to colorless (post-enrichment) (Fig. 4b). Flow cytometry analysis further demonstrated that the proportion of GalOx-*E. coli* cells increased to 62%, 87%, and 92% from initial populations of 1%, 5% and 10%, respectively, after the magnetic separation (Fig. 4c). These results confirm the capability of the GENie platform to efficiently enrich active oxidase-expressing *E. coli* cells from heterogeneous populations, supporting its application in practical directed evolution campaigns.

**Figure 4.**
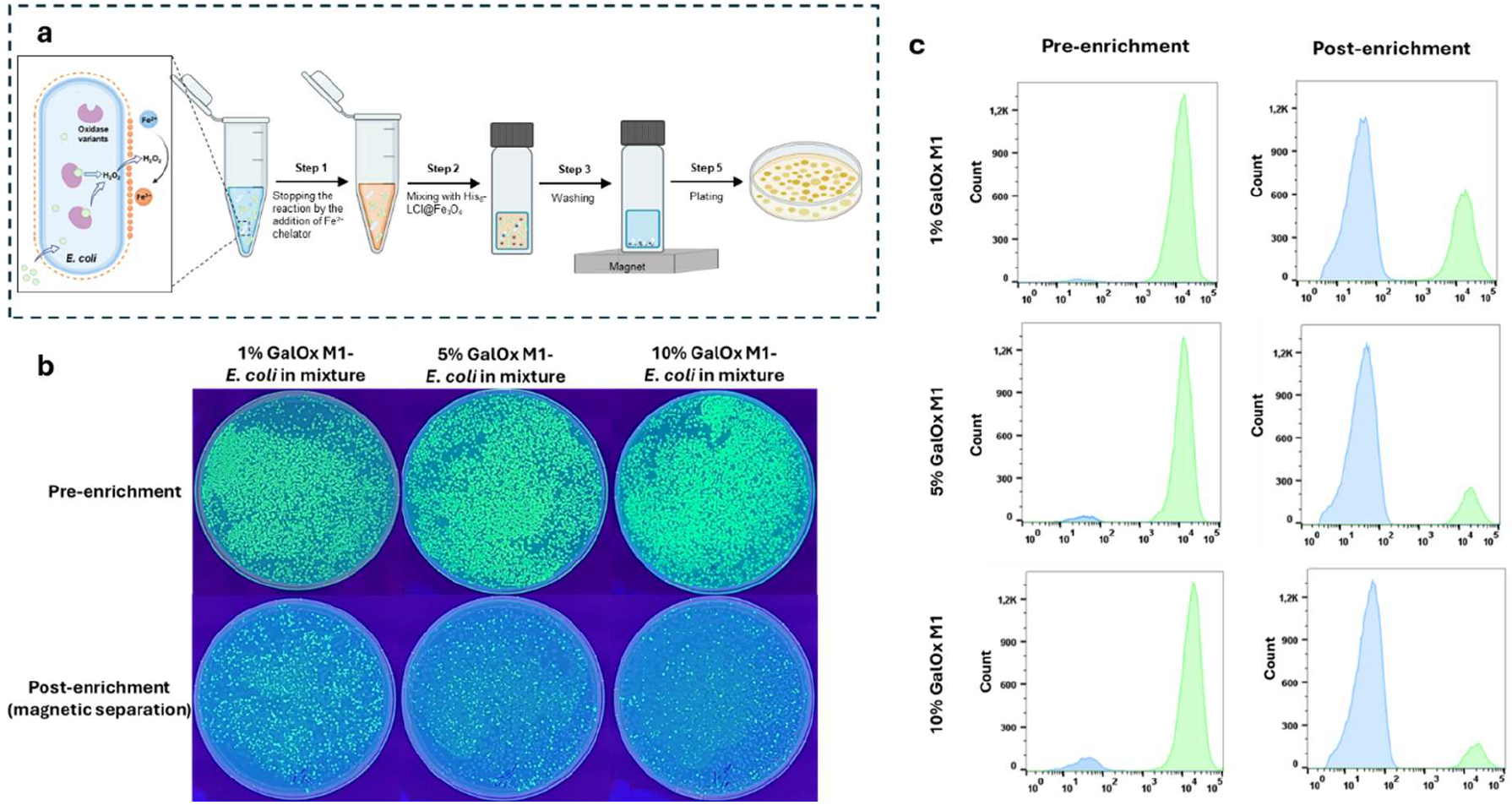
Proof of concept of the GENie platform for effective enrichment of target GalOx M1-E. *coli* cells from a cell mixture. (a) Cell mixtures are incubated with His_6__LCI@Fe_3_O_4_ beads, following the magnetic separation. (b) The auto-induction agar plates enable the quick visualization of the enrichment of target GalOx M1-E. *coli* cells before and after magnetic separation. The first row shows plates with initial GalOx M1-*E. coli* cell ratios before enrichment. In the second row, plates display the GalOx M1-*E. coli* and eGFP-*E. coli* colonies distribution after enrichment. (c) Flow cytometry for the statistical evaluation of the GalOx M1-*E. coli* cells and eGFP-*E. coli* cells before and after magnetic separation (the total number of events is 150,000).

### Directed evolution of FgGalOx

Then, we applied the GENie platform to directed enzyme evolution campaigns, selecting galactose oxidase from *Fusarium graminearum* NRRL 2903 (FgGalOx) as a model enzyme to evaluate its performance(*27, 28*). We generated a random mutagenesis library of GalOx using error-prone PCR (epPCR) followed by Megawhop cloning, yielding a library size exceeding 10^7^ transformants. During the evolution of GalOx, we performed two rounds of enrichment. In the second round, two distinct selection pressures—reduced substrate concentration and shortened reaction time—were applied to identify variants with improved substrate affinity and enhanced catalytic turnover, respectively. After magnetic separation, we recovered the collected populations on agar plates (Fig. S6) and re-assessed their activity profiles by the ABTS assay. Fig. 5a illustrates the workflow of directed enzyme evolution using the GENie platform. For the GalOx library, we evaluated activity improvements and quantified the number of enhanced variants in each round, both before and after magnetic enrichment. A substantial increase in the number of improved variants was observed after enrichment in each round of screening (Fig. 5b). In the first round, we identified 37 variants exhibiting ≥ 1.5-fold improved activity after magnetic enrichment (out of 96 screened variants); in contrast, 3 variants showed this improvement before enrichment. In the second round, we identified 30 and 25 improved variants exhibiting ≥ 1.5-fold improved activity under the two selection pressures, respectively, whereas no variants showed this improvement before magnetic enrichment (Fig. 5c). Among the beneficial variants identified in the first round, the top-performing GalOx variant, G1 (M1 + T220I/C383S/N413D), exhibited a 6.1-fold increase in *k*_cat_/*K*_m_ over its parent and an 11.3-fold increase in *k*_cat_/*K*_m_ over the wild type. Following the second round of screening, two additional superior GalOx variants were identified, demonstrating up to a 26-fold improvement in catalytic efficiency compared to the wildtype. Notably, the variant G2(*K*_m_) (G1 + S20C/I498T/N521S/N537K) showed a 5.8-fold decrease in *K*_m_ (57 mM to 9.9 mM), indicating markedly improved substrate affinity driven by the low-substrate-concentration screening pressure. In contrast, variant G2(*k*_cat_) (G1 + Q22R/F295L/N298S) exhibited a 7.8-fold increase in *k*_cat_ (1070 s^−1^ to 8349 s^−1^), reflecting a substantially enhanced turnover rate, which resulted from the screening pressure imposed by shortened reaction time (Fig. 5d, 5g, and S7). Notably, except I498T, all identified mutations are distal to the active site (Fig. 5e and 5f). These results demonstrate that the GENie platform not only increases the recovery rate of improved variants but also enables targeted evolution of distinct kinetic properties under defined selection pressures. The substantial gains in catalytic efficiency across successive rounds highlight the system’s capacity to tailor enzyme properties in a controllable manner and to explore long-range effects in enzymatic catalysis.

**Figure 5.**
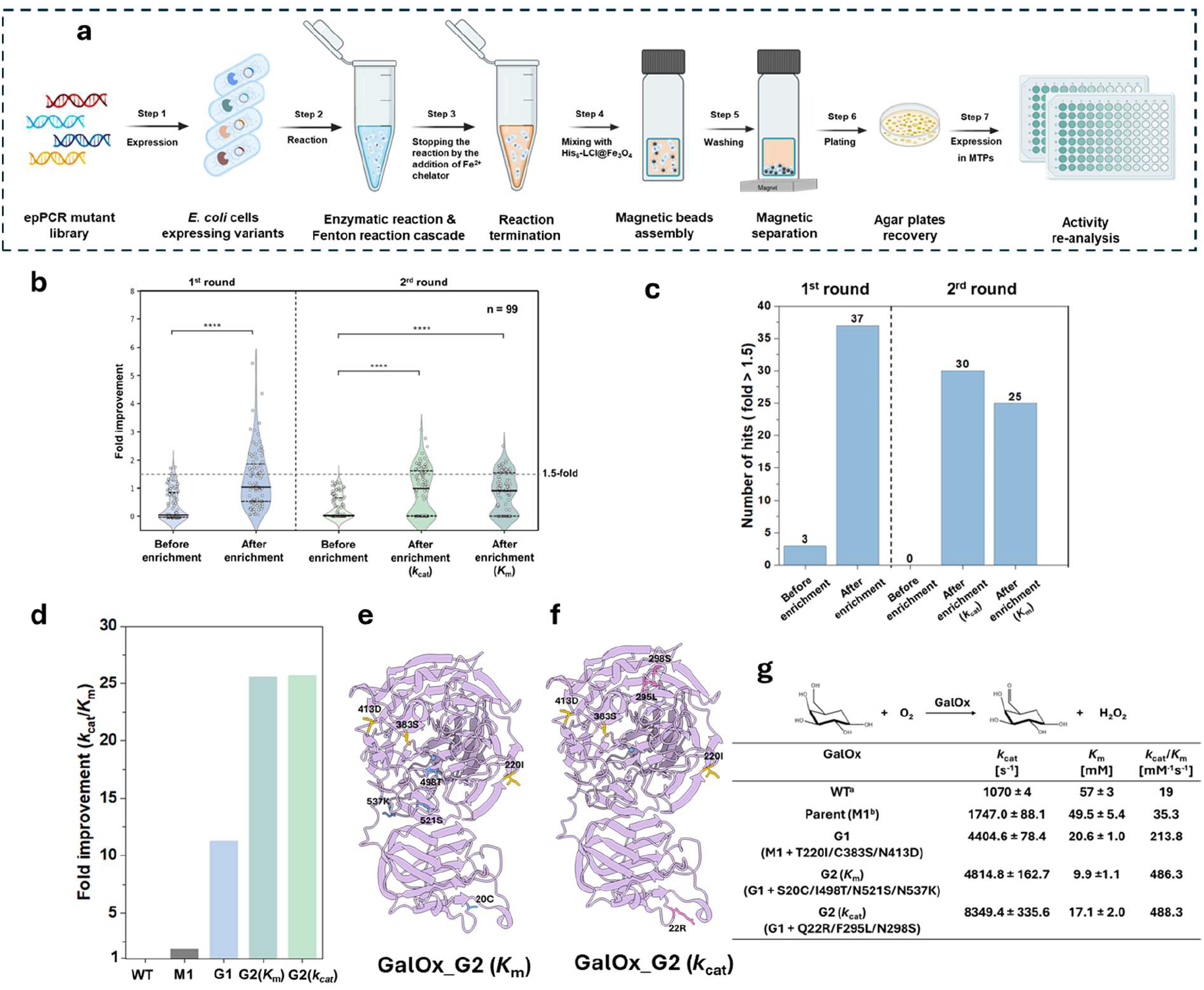
(a) Workflow of the magnetic-based screening platform. (b) Enrichment outputs of the two rounds of magnetic separation before and after sorting. (c) The number of improved hits in the two rounds of magnetic separation before and after sorting. The numbers are calculated from one micro-tiler plate (96 events). (d) Fold improvement of the best variants of GalOx. (e-f) Mapping of the mutations on the structure of GalOx (PDB: 2EIE). (g): Kinetic characterization of the best variants for GalOx. [a] Data from Sun et al., 2001(*27*). [b] GalOx M1 contains five mutations: S10P/M70V/G195E/V494A/N535D.

### Directed evolution of FgGalOx, GtAOX, and RgDAAO

Having successfully applied the magnet-based screening system to the directed evolution of GalOx, we then extended the platform to amino acid oxidase and alcohol oxidase to validate its general applicability. For both enzymes, we generated random mutagenesis libraries comprising more than 10^7^ transformants. In the case of alcohol oxidase, we targeted the alcohol oxidase from *Gloeophyllum trabeum* (GtAOx) to improve its catalytic efficiency toward the non-natural substrate benzyl alcohol, a model reaction relevant to pharmaceutical and fine chemical synthesis(*29*). After a single round of magnetic screening and enrichment, we identified the best-performing AOx variant among 6 candidates (Fig. 6a and S8), AOx_V6 (F101S/G254R/D544N), exhibiting a 25-fold increase in specific activity compared to the WT (Fig. 6b and 6c). Notably, the *K*_m_ value of AOx_V6 showed a 12-fold decrease, from 90.2 mM to 7.5 mM, indicating a significantly enhanced substrate affinity (Fig. 6d, S9 and S10). Given the negligible activity of wild-type GtAOx toward aromatic alcohols, the significant decrease in *K*_m_ coupled with the pronounced activity enhancement underscores the power of the GENie platform to rapidly enrich and identify rare, high-performance variants, ultimately enabling effective functional reprogramming of the enzyme toward non-natural substrates.

**Figure 6.**
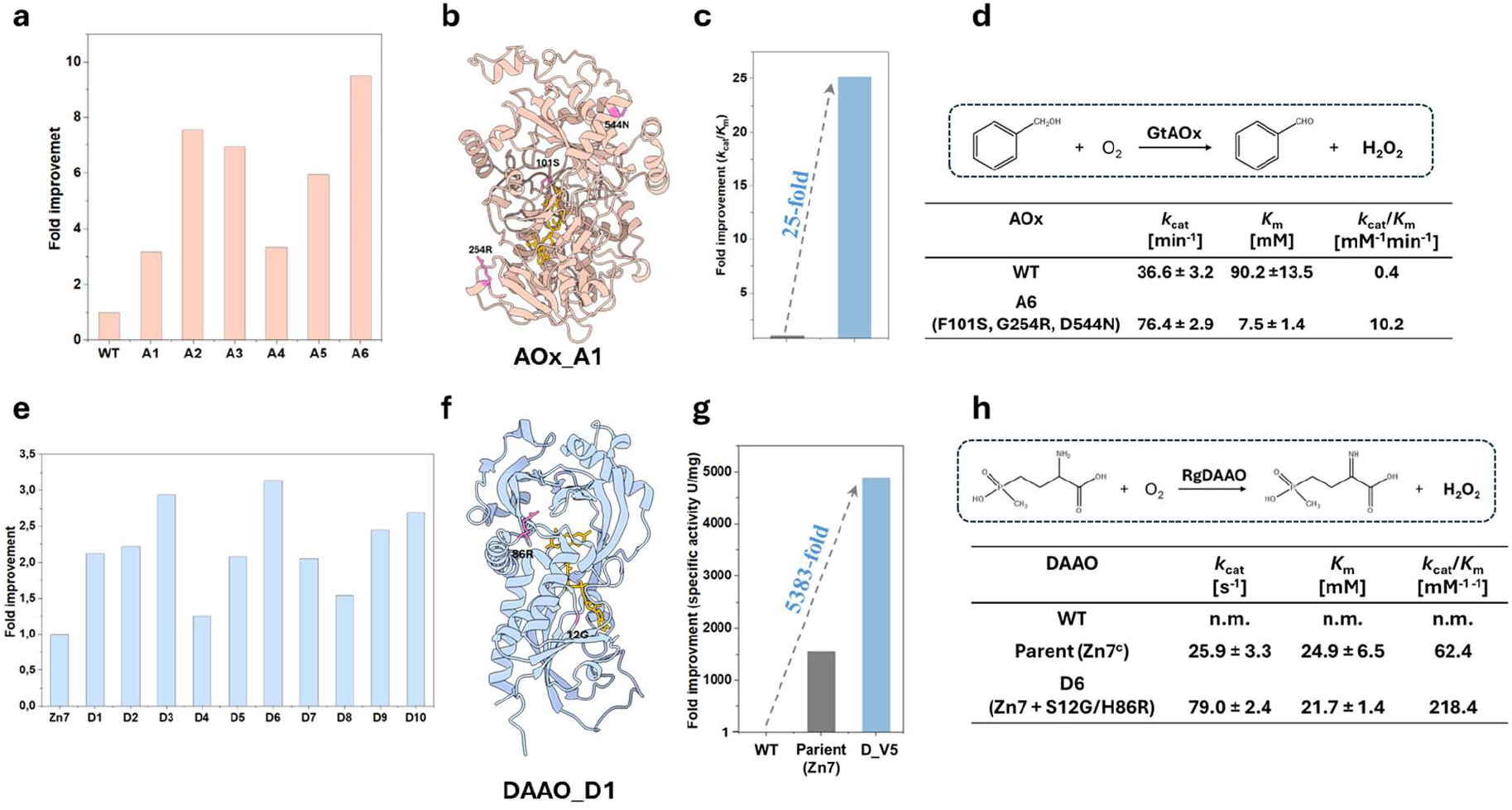
(a) The improvement fold of the 6 best-performing AOx variants after enrichment measured by GC. (b) Mapping of the mutations on the AlphaFold 3 model of AOx. (c) Fold improvement of the best variants of AOx. (d) Kinetic characterization of the best variants for AOx. (e) The improvement fold of the 10 best-performing DAAO variants after enrichment measured by absorbance (420nm). (f) Mapping of the mutations on the structure of DAAO (PDB: 1C0K). (g) Fold improvement of the best variants of DAAO calculated from specific activity. (h) Kinetic characterization of the best variants for DAAO. DAAO Zn7 contains three mutations: N54T/M213T/S335Q^[26]^.

We further target the D-amino acid oxidase from *Rhodotorula gracilis* (RgDAAO) to identify the variants with improved catalytic activity toward the non-natural substrate D-glufosinate, which can be converted into the corresponding keto acid, 2-oxo-4-[(hydroxy) (methyl) phosphinyl] butyric acid (PPO), a precursor of an important herbicide(*30–32*). Following a single round of magnetic screening and enrichment, we identified the best-performing DAAO variant among 10 candidates (Fig. 6e and S11), DAAO_D6 (M1 + S12G/H86R, 118.70 U mg^−1^), exhibiting a 3.4-fold increase in specific activity over the parent (35.16 U mg^−1^) and a 5383-fold increase over WT (0.02 U mg^−1^) (Fig. 6f and 6g). Due to the extremely low activity of the WT toward D-glufosinate, its kinetic parameters could not be reliably determined. Notably, the introduction of the distal double mutation S12G and H86R through random mutagenesis further enhanced catalytic activity by 3.5-fold relative to the parent (DAAO Zn7)(Fig. 6h and S12), despite the activity center of the variant Zn7 having been saturated and extensively studied in previous research(*33*). These results highlight the power of the GENie platform in overcoming apparent evolutionary plateaus and uncovering hidden fitness gains that may not be predictable by rational design alone.

## Discussion

Directed evolution has long been constrained by complex screening hardware and labor-intensive workflows. By transducing enzymatic turnover into a magnetically capturable property, we decoupled uHTS from specialized instrumentation and established a first genuine test-tube screening platform that streamlines protein engineering workflows. Beyond its ultra-high screening capacity, this approach significantly lowers the barrier to entry for directed evolution, transforming an infrastructure-intensive process into a benchtop approach that requires minimal equipment, thereby broadening the accessibility of directed evolution across diverse laboratory settings and its applications in synthetic biology.

The robustness and general applicability of the genuine *test-tube* screening platform are validated by the substantial evolutionary gains across multiple enzymes. Using GalOx as a model, the system efficiently navigated the fitness landscape under different selection pressures, yielding variants with up to 26-fold improvements in catalytic efficiency after two rounds of screening. Notably, the platform steered enzyme evolution under defined selection pressures, identifying a variant with a 5.8-fold decrease in *K*_m_ and a variant with a 7.8-fold increase in *k*_cat,_ demonstrating its capability for rapid and controllable optimization of specific enzyme properties. The power of the GENie platform was further highlighted when applied to D-AAO and AOx. In traditional protein engineering, finding a modest performance hit often demands multiple rounds of screening(*34*). The GENie platform isolated D-AAO and AOx variants with up to 5383-fold and 25-fold improvements over their respective wild types after a single round of screening. Together, these sweeping improvements suggest that this hardware-free platform could markedly accelerate the engineering of any H_2_O_2_-generating oxidase, which is a class of biocatalysts essential to the future of pharmaceuticals, biosensors, and green chemistry.

The genuine test-tube screening platform circumvents the fundamental physical and operational limitations that restrict current uHTS technologies. In the genuine test-tube screening platform, a physical genotype–phenotype linkage is established through His_6_-tagged peptide–functionalized magnetic beads and localized deposition of Fe^3+^ on the surfaces of catalytically active cells, achieving a screening throughput of >10^8^ events s^−1^ and an enrichment factor of up to 63-fold. This spatial physical capture of Fe^3+^ on the cell surface avoids the droplet effects and diffusion-mediated crosstalk inherent to flow cytometry, achieving effective enrichment without the need for physical compartmentalization. Moreover, both flow cytometry and microfluidic screening systems are inherently serial processes, analyzing individual cells or droplets one by one, thereby constraining overall throughput by event and sorting rates. In addition, these uHTS systems typically rely on customized fluorogenic substrates, which can lead to enzymes optimized for artificial probes rather than their true catalytic substrates(*35–37*). By transitioning from serial interrogation to bulk magnetic separation, the platform enables true parallel enrichment of large variant populations. The GENie platform simplifies the traditionally complex workflows of uHTS, which encompasses multi-step labeling, compartmentalization, and sorting procedures, into a streamlined process involving single-pot incubation followed by rapid magnetic capture, while directly selecting enzymes against their intended catalytic substrates. As a result, one round of directed enzyme evolution can be completed within less than one week. The reduced complexity lowers costs, shortens experimental timelines, and minimizes instrumentation requirements, while compatibility with standard liquid-handling automation for scalable processing and data generation. A detailed comparison with existing uHTS technologies is shown in Tab. 1.

**Table 1:** Comparison of the current uHTS technology.

| Parameter | Magnetic bead-based screening | Flow cytometry-based screening | Microfluidic droplet screening |
| --- | --- | --- | --- |
| **Screening principle** | Bulk enrichment via magnetic separation | Fluorescence detection and sorting of individual cells | Compartmentalized reactions in droplets |
| **Screening format** | Parallel bulk selection | Serial single-cell analysis | Serial droplet analysis |
| **Maximum library size** | $\sim 10^9$–$10^{12}$ variants | $\sim 10^7$–$10^8$ variants | $\sim 10^6$–$10^8$ variants |
| **Throughput limitation** | Not limited by the detection rate | Limited by the flow cytometer event rate | Limited by droplet generation and sorting rate |
| **Experimental workflow** | Simple (incubation → magnetic separation) | Multi-step labelling and sorting | Droplet generation, incubation, and sorting |
| **Instrumentation requirement** | Minimal (magnetic rack) | Specialized flow cytometer | Microfluidic chip and droplet sorter |
| **Equipment cost** | Low | High | High |
| **Automation compatibility** | High (simple liquid handling) | Moderate | Technically complex |
| **Large Data set acquisition** | Easy | Difficult | Difficult |
| **Key advantage** | Parallel library screening with a simple workflow | Single-cell resolution | Compartmentalized activity assays |

We envision that the genuine test-tube screening platform will enable more researchers across diverse laboratories to perform directed evolution of oxidases and other enzymes (e.g., by enabling cascade reactions within cells). Additionally, this platform will also enhance AI and machine learning applications by rapidly expanding datasets of high-performance oxidase variants(*38*). In enzyme science, such extensive and high-quality datasets provide a critical foundation for developing accurate predictive models, thereby significantly accelerating enzyme activity prediction, functional annotation, and pathway optimization(*39–41*).

## MATERIALS AND METHODS

### Materials

Fe_3_O_4_ nanoparticles in water dispersion were purchased from PlasmaChem GmbH (Germany). Glass vials (1.5 mL) were obtained from Th. Geyer GmbH (Germany). Taq DNA polymerase was purchased from New England Biolabs (Frankfurt, Germany). Plasmid extraction and PCR purification kits were purchased from Macherey-Nagel (Düren, Germany). Microtiter plates (flat-bottom transparent MTPs, and v-bottom transparent MTPs) were purchased from Greiner Bio-One (Frickenhausen, Germany). Glufosinate-ammonium was ordered from TCI Deutschland GmbH (Germany). Galactose, benzyl alcohol, benzaldehyde, HRP, ABTS, iron (II) chloride, iron (III) chloride, and other laboratory-grade chemicals were purchased from Sigma-Aldrich (Merck, Germany) or AppliChem (Germany) unless specified.

### Expression and purification of His-DZ-LCI and eGFP-MBP1

The DNA sequences encoding His-DZ-LCI and eGFP-MBP1 were synthesized and cloned into the pET-28a(+) vector by GenScript Biotech (Netherlands). The resulting plasmids were transformed into competent E. coli BL21(DE3) cells for protein expression. Following overnight incubation at 37 °C in a shaker, the pre-culture was used to inoculate 50 mL of Lysogeny Broth (LB) medium (supplemented with 50 µg/mL kanamycin) and cultured at 37 °C with continuous shaking at 200 rpm. When the OD600 reached 0.4-0.6, the main cultures were induced with 0.1 mM IPTG and further incubated at 18 °C, 200 rpm overnight. Cells were harvested by centrifugation, resuspended in lysis buffer (50 mM Tris-HCl, pH 8, 300 mM NaCl), and lysed by sonication. The clarified lysate obtained after centrifugation was filtered through a 0.45 μm membrane and applied to a Ni-NTA Sepharose column (1 mL, MACHEREY-NAGEL GmbH) pre-equilibrated with lysis buffer. The column was washed sequentially with lysis buffer containing 0, 10, and 20 mM imidazole (5 mL each), and proteins were eluted with lysis buffer containing 250 mM imidazole. The eluted proteins were buffer-exchanged into the final storage buffers (50 mM Tris-HCl, pH 8 for His-DZ-LCI; 50 mM Tris-HCl, pH 8, 300 mM NaCl for eGFP-MBP1) using a PD-10 desalting column (Cytiva). All purification steps were performed at 4 °C, and protein concentrations were determined by UV absorbance at 280 nm.

### Functionalization of iron oxide beads

The purified protein (5 mM His-DZ-LCI) was incubated with Fe_3_O_4_ nanoparticles (0.75 mg/L) in functionalization buffer (50 mM Tris-HCl, pH 8) in a total volume of 200 µL, with shaking at 1200 rpm in glass vials (Screw neck vial ND 8, TH. GEYER GmbH & Co. KG, Germany) for 3 min. The functionalized nanoparticles were then collected by magnetic separation, washed with water, and finally resuspended in water for subsequent use.

### Expression of mCherry, eGFP, and GalOx

The DNA sequences encoding mCherry, eGFP, and GalOxM1 were synthesized and cloned into the pET-28a(+) vector by GenScript Biotech (Netherlands). The resulting plasmids were transformed into competent *E. coli* BL21(DE3) cells for protein expression. After overnight cultivation at 37 °C with shaking, the pre-culture was used to inoculate 50 mL of LB medium supplemented with 50 µg/mL kanamycin. Cultures were grown at 37 °C with shaking at 200 rpm until the OD_600_ reached 0.4–0.6, at which point protein expression was induced with 0.1 mM IPTG for mCherry and eGFP, and with 0.2 mM IPTG and 0.4 mM Cu^2+^ for GalOx. The cultures were then incubated at 18 °C with shaking at 200 rpm overnight for protein expression.

### Enrichment assay developed by mCherry and eGFP

Target cells (*E. coli* expressing mCherry intracellularly) were first modified with Fe^3+^ on the cell surface by incubating cell suspensions (OD_600_ = 4 in 0.9% (w/v) NaCl) with 100 μM FeCl_3_ for 5 min at room temperature (total volume 100 µL). The cells were then washed with 0.9% NaCl and resuspended in 100 µL of the same solution. Cell mixtures containing 10%, 5%, or 1% Fe^3+^-modified mCherry-expressing *E. coli* (with the remainder being unmodified *E. coli* expressing eGFP) were prepared at a final OD_600_ of 0.1.

Glass vials were pre-blocked by incubation with 500 µL of 5 µM eGFP-MBP1 in 50 mM Tris-HCl (pH 8) at 200 rpm for 5 min, followed by washing with Tris-HCl buffer and 0.9% NaCl. Iron oxide bead functionalization was carried out in these pre-blocked vials. After washing and resuspension in water, the beads were transferred to a new pre-blocked vial, and residual liquid was removed by magnetic separation. The prepared cell mixtures (200 µL) were then added to the beads and incubated with shaking at 1200 rpm for 5 min. Washing was performed by magnetic separation (2 min), removal of the supernatant, resuspension in 200 µL NaCl, and 3 min water bath sonication; this process was repeated three times. Finally, the cell–bead complexes were resuspended in 200 µL NaCl and plated onto auto-induction agar plates.

### Enrichment assay developed by GalOx and eGFP

Cell mixtures containing 10%, 5%, or 1% GalOx-expressing *E. coli* (with the remainder being *E. coli* expressing eGFP) were prepared at a final OD_600_ of 0.5 in 0.9% NaCl and incubated with 300 µM Fe^2+^. A 200 µL aliquot of this mixture was then added to 800 µL 0.9% NaCl containing 10 mM galactose (final concentration). After immediate vortexing, the suspension was incubated at room temperature for 20 min to allow the oxidation reaction and generation of Fe^3+^ on the surface of GalOx-expressing cells. The reaction was terminated by adding 18 µL of 10 mM 1,10-Phenanthroline. The resulting mixture was subsequently added to the functionalized magnetic beads for the enrichment assay as described above.

### Error-prone PCR mutagenesis and cloning of the mutated gene library

For epPCR library generation, the gene fragments encoding GalOx, RgDAAO, and GtAOX were amplified using designed primers and Taq polymerase under the PCR conditions listed in Tab. S1, S2, and S3. The PCR products were analyzed by agarose gel electrophoresis, and bands of the expected size were excised and purified. The resulting variant libraries were cloned into pET-28a(+) using the MEGAWHOP method (Tab. S4 and S5). PCR products were treated with DpnI (20 U, 37 °C, 2 h) to remove template DNA and subsequently purified. For library construction, 100 ng of each purified MEGAWHOP product was transformed into 25 µL of NEB 10-beta electrocompetent *E. coli* cells (New England Biolabs, Frankfurt, Germany); based on our estimate, at least 8 transformations yield a library of ~10^7^ variants. After recovery in SOC medium at 37 °C for 1 h, cells were directly inoculated into LB medium (5 mL per transformation) and cultured overnight at 37 °C. Plasmids were then extracted, and 100 ng of each purified library plasmid was transformed into 25 µL of self-prepared BL21(DE3) electrocompetent *E. coli* cells for protein expression.

### Expression and purification of FgGalOx, RgDAAO, and GtAOX

The DNA sequences encoding RgDAAO (Zn7) and GtAOX were synthesized and cloned into the pET-28a(+) vector by GenScript Biotech (Netherlands). The resulting plasmids were transformed into competent *E. coli* BL21(DE3) cells for protein expression. Following overnight incubation at 37 °C in a shaker, the pre-culture was used to inoculate 50 mL of LB medium (supplemented with 50 µg/mL kanamycin) and cultured at 37 °C with continuous shaking at 200 rpm. When the OD_600_ reached 0.4-0.6, the main cultures were induced by adding 1% lactose (m/v) at 25°C for RgDAAO, and by adding 0.2 mM IPTG at 18 °C for GtAOX, shaken at 200 rpm overnight. Cells were harvested by centrifugation, resuspended in lysis buffer (50 mM Tris-HCl, pH 8 for Rg DAAO; 100 mM NaPi, pH 7 for GalOx and GtAOX), and lysed by sonication. The clarified lysate obtained after centrifugation was filtered through a 0.45 μm membrane and applied to a Ni-NTA Sepharose column (1 mL, MACHEREY-NAGEL GmbH) pre-equilibrated with lysis buffer. The column was washed sequentially with lysis buffer containing 0, 10, and 20 mM imidazole (5 mL each), and proteins were eluted with lysis buffer containing 250 mM imidazole. The eluted proteins were buffer-exchanged into the final storage buffers (same as lysis buffer) using a PD-10 desalting column (Cytiva). All purification steps were performed at 4 °C, and protein concentrations were determined by UV absorbance at 280 nm for GalOx, and by BCA assay for RgDAAO and GtAOX.

### FgGalOx, RgDAAO, and GtAOX epPCR library pre-screening by magnetic enrichment

Enzyme-expressing E. coli cells (OD_600_ = 0.5) in 0.9% NaCl containing 300 µM Fe^2+^ were incubated with substrate at defined concentrations (10 mM galactose for GalOx, 50 mM glufosinate-ammonium for RgDAAO Zn7, and 0.25 mM benzyl alcohol for GtAOX). A series of reactions with varying incubation times was performed, followed by centrifugation at 11,000 rpm for 30 s. Formation of a compact, pointed cell pellet was used as an indicator that sufficient Fe^3+^ had been generated for cell surface decoration. For screening improved variants, substrate concentration can be reduced to select for lower Km, or reaction time shortened to select for higher kcat. Once optimal conditions were established, cells were incubated with Fe^2+^ and substrate under defined conditions, and the reaction was terminated by adding 1,10-phenanthroline (final concentration 10 mM). The resulting mixture was then subjected to magnetic bead–based enrichment as described above.

### Protein expression and preparation of crude cell extracts in 96-well microtiter plates (MTPs)

Single colonies were picked and cultured in 96-well microtiter plates (MTPs, flat-bottom, transparent) containing 150 µL LB medium supplemented with kanamycin at 37 °C overnight (5–10 plates were sufficient to identify improved variants). Subsequently, 5 µL of each culture was transferred into new 96-well MTPs (V-bottom, transparent) containing 135 µL LB medium and incubated at 37 °C for 1.5 h. Protein expression was then induced according to the oxidase induction conditions used in flask cultures. After cultivation, cells were harvested by centrifugation, and the resulting pellets were frozen at ™80 °C for 30 min. Cell lysis was performed by adding 100 µL lysozyme solution (1.5 mg/mL) to each well, followed by resuspension and incubation at 37 °C for 30 min. After a second centrifugation step, the clarified supernatant was collected and used for subsequent activity assays.

### FgGalOx, RgDAAO, and GtAOX epPCR library rescreening by colormetric assay

Hydrogen peroxide (H_2_O_2_) generated by oxidases during substrate oxidation was monitored using horseradish peroxidase (HRP)-coupled colorimetric assays. For GalOx and RgDAAO, activity was determined by adding 2 mM ABTS and 5 U/mL HRP in NaPi buffer (100 mM, pH 7.0) for GalOx or Tris buffer (50 mM, pH 8.0) for RgDAAO, together with the respective substrate (galactose for GalOx and glufosinate-ammonium for RgDAAO). The reaction was initiated by adding 20 µL of cell lysate to 180 µL of the ABTS assay mixture in 96-well microtiter plates (flat-bottom, transparent), and the increase in absorbance was monitored at 420 nm using a Tecan Sunrise™ plate reader. For GtAOX, activity was measured using a colorimetric assay containing 0.1 mM 4-aminoantipyrine, 1 mM dichloro-2-hydroxybenzenesulfonic acid, and 5 U/mL HRP in NaPi buffer (100 mM, pH 7.0), together with the substrate benzyl alcohol. The formation of a pink-purple quinone product was monitored by measuring absorbance at 515 nm using the same plate reader.

### Kinetic characterization of the purified FgGalOx, RgDAAO, and GtAOX variants

Top candidates identified from rescreening were selected for expression in flasks and purified using the Ni-affinity column. For FgGalOx and RgDAAO, initial velocities were determined using the ABTS assay. Reactions (200 µL) containing varying substrate concentrations (galactose: 0.1–125 mM for GalOx; glufosinate-ammonium: 0.1–50 mM for RgDAAO), buffer (100 mM NaPi, pH 7.0 for GalOx; 50 mM Tris, pH 8.0 for RgDAAO), and enzyme (0.012 nM for GalOx and 6.5 nM for RgDAAO) were carried out in 96-well plates (flat-bottom, transparent). The increase in absorbance at 420 nm was monitored at 25 °C for 10 min using a Tecan Sunrise™ plate reader.

For GtAOX, kinetic parameters were determined by GC analysis. Reaction mixtures (200 µL) containing benzyl alcohol (0.5–100 mM), buffer (100 mM NaPi, pH 7.0), and enzyme (26 µM for wild type and 6 µM for variant) were incubated at 25 °C with shaking (600 rpm) for 10 min in 1.5 mL tubes. Reactions were quenched with 10 µL of 6 M HCl, followed by extraction with 200 µL ethyl acetate. After centrifugation, the organic phase was transferred to glass vials for GC analysis (Shimadzu GC-2010, FID). Separation was performed using a HYDRODEX BETA-TBDAc column (MACHEREY-NAGEL GmbH, Germany) with an inlet temperature of 280 °C, detector temperature of 280 °C, initial oven temperature of 50 °C (2 min), followed by a ramp of 10 °C·min^−1^ to 200 °C (10 min hold). Quantification was based on calibration curves prepared from authentic standards unless otherwise stated.

## Supporting information

GENie_support_Final

## Acknowledgments

We acknowledge Petra Esser, Stefan Hauk, and Dr. Rostislav Vinokur from DWI—Leibniz-Institut für Interaktive Materialien for their help with materials characterization. We acknowledge ChatGPT for polishing manuscript writing. Some of the figures were created using https://Biorender.com. Protein molecular graphics were performed with UCSF ChimeraX 1.4, developed by the Resource for Biocomputing, Visualization, and Informatics at the University of California, San Francisco, with support from National Institutes of Health R01-GM129325 and the Office of Cyber Infrastructure and Computational Biology, National Institute of Allergy and Infectious Diseases.

## Author contributions

Conceptualization: L.L.F., M.M.C., U.S.; Methodology: L.L.F., M.M.C.; Investigation: L.L.F., M.M.C.; Visualization: L.L.F., M.M.C.; Funding acquisition: U.S.; Supervision: U.S.; Writing – original draft: L.L.F., M.M.C.; Writing – review and editing: L.L.F., M.M.C., U.S.

## Data and materials availability

All data needed to evaluate the conclusions in this paper are present in the paper and/or the Supplementary Materials.

