## Supplementary material for "Genuine Directed Evolution In Test Tube (GENie)": GENie_support_Final

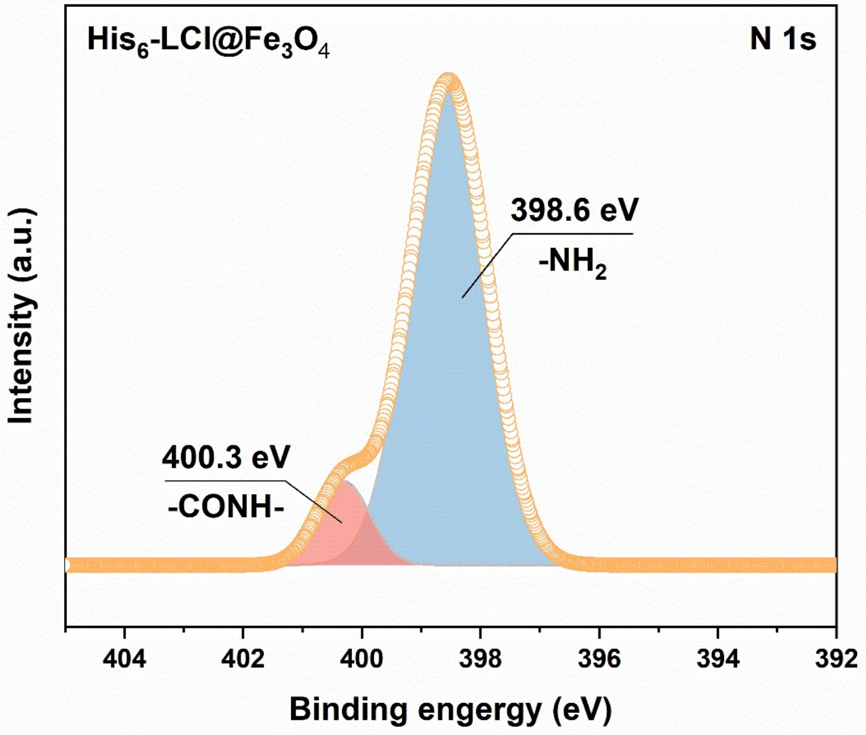


**Figure S1. XPS spectra (N 1s) of His$_6$-LCl@$Fe_3O_4$.** The presence of peptide-associated -NH2 and -COHN- groups indicates the successful modification.

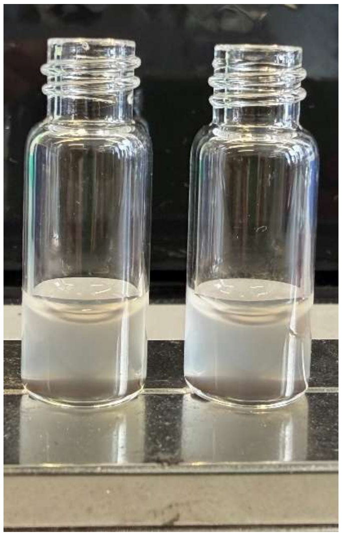

**Figure S2. Imaging of $Fe^{3+}$-deposited *E. coli* cells captured by peptide-modified iron oxide beads without his-tag and bare *E. coli* cells captured by his-tagged peptide-modified iron oxide beads.** A cloudy supernatant after magnetic extraction suggests that magnetic capture is initiated only through interactions between $Fe^{3+}$ on the cell membrane and the His-tag on the bead surface.

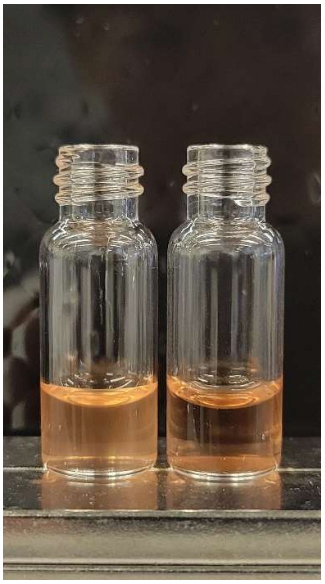

**Figure S3. Imaging of $Fe^{3+}$-deposited *E. coli* cells before and after magnetic extraction with the presence of 1,10-phenanthroline.** A clear supernatant after magnetic extraction indicates that 1,10-phenanthroline does not interfere with the magnetic extraction.

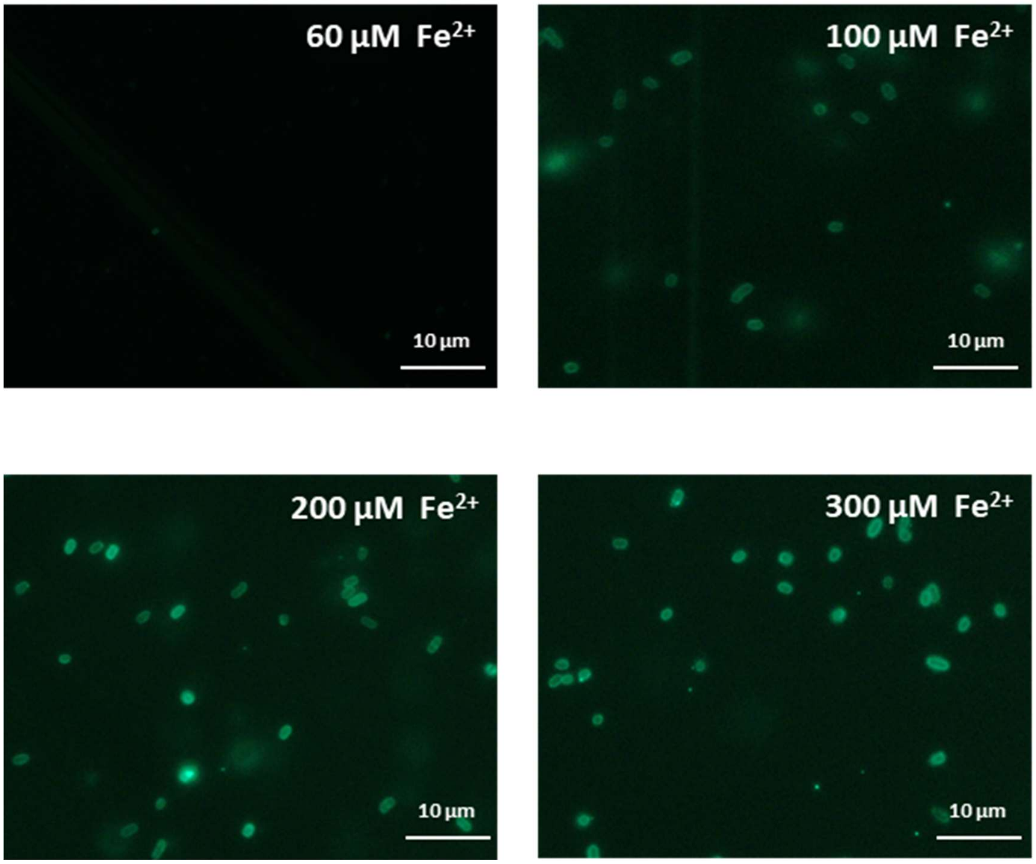


**Figure S4. Fluorescence imaging of *E. coli* cells incubated with varying concentrations of $Fe^{2+}$, followed by labeling with His-eGFP.** When the $Fe^{2+}$ concentration falls below 60 µM for OD 0.1 *E. coli* cells, no background is observed.

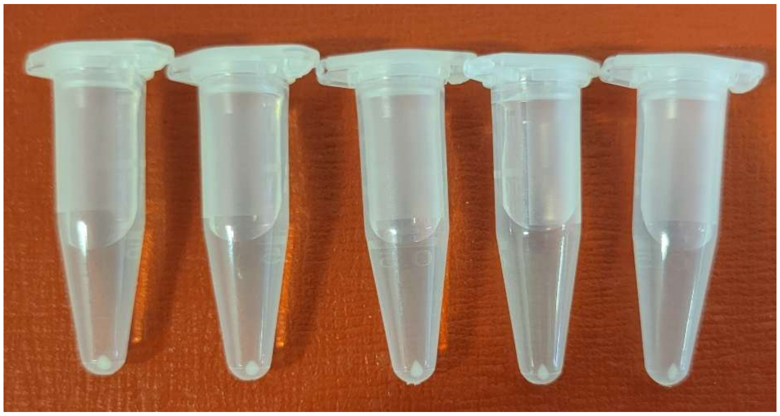

**Figure S5. Imaging of GalOx-expressing *E. coli* pellets after varying reaction times in the presence of $Fe^{2+}$.** From left to right, reaction times are 10, 15, 20, 25, 30 min, with 10 mM galactose. As time progresses, the cell pellets become increasingly pointed.

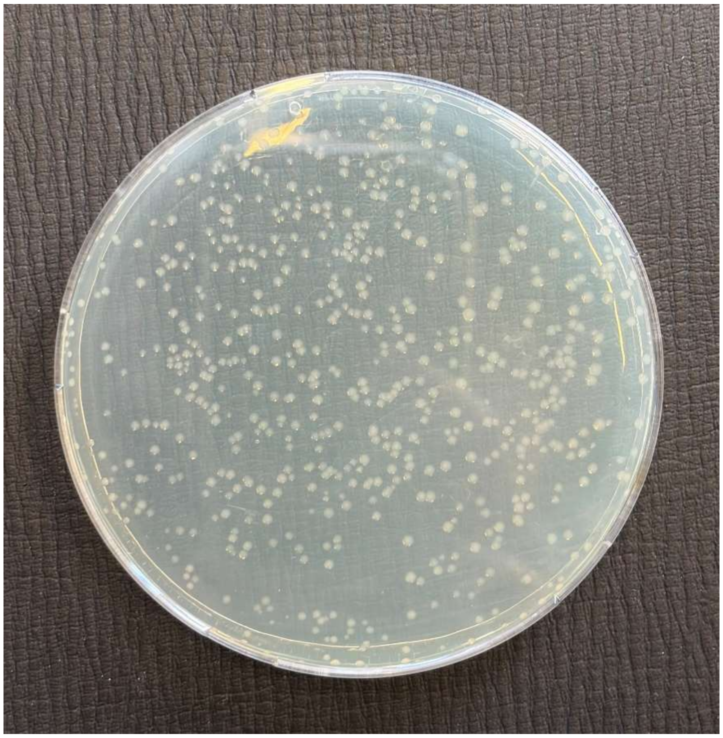

**Figure S6. Imaging of *E. coli* colonies growing on an agar plate after uHTS by magnetic extraction.** Following magnetic extraction of cells from the library (OD 0.1), approximately 500–1000 colonies were observed on the agar plates.

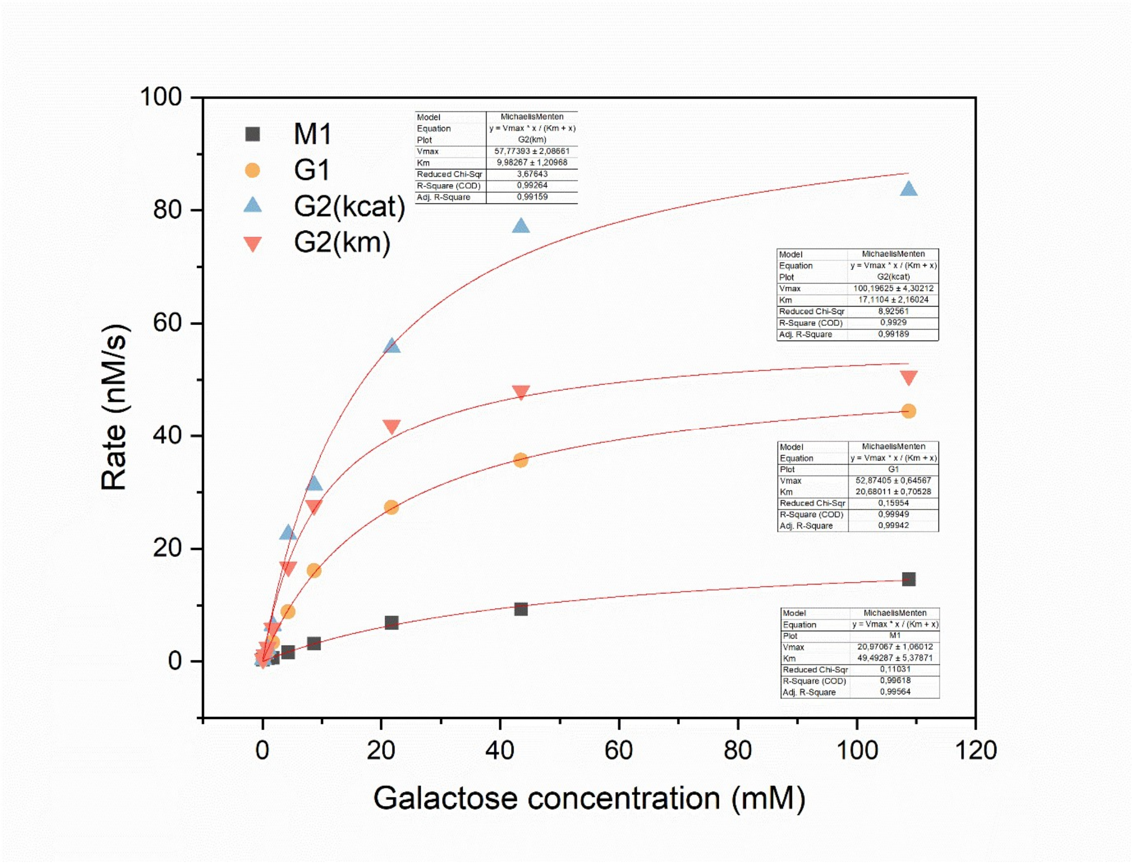


**Figure S7. Kinetics curve of Galactose oxidases fitted with Michaelis-Menten.**

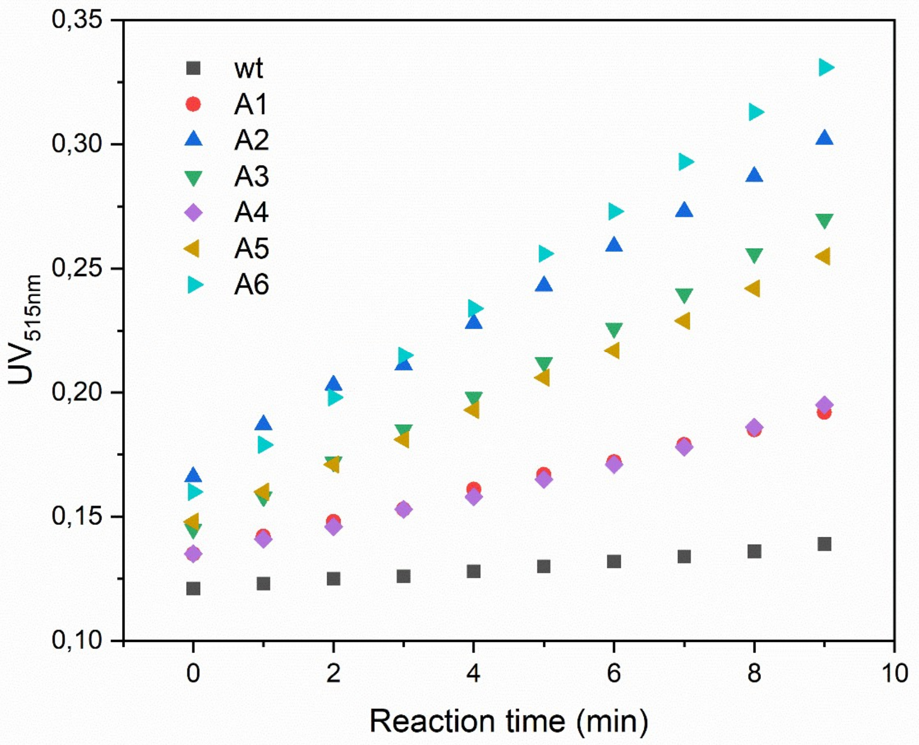


**Figure S8. Kinetics curve of purified GtAOx variant candidates.** Six improved GtAOx candidates are identified after a single round of screening.

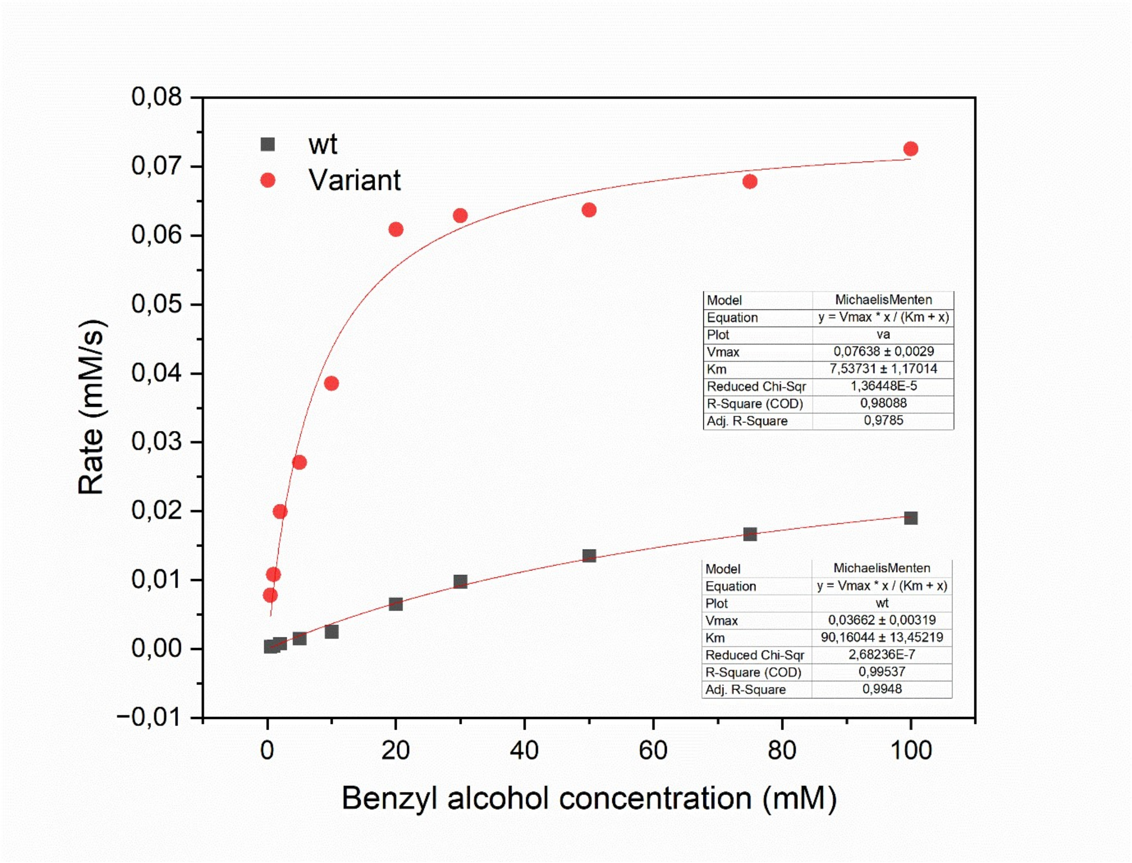


**Figure S9. Kinetics curve of GtAOx (wild type and variant A6) fitted with Michaelis-Menten.**

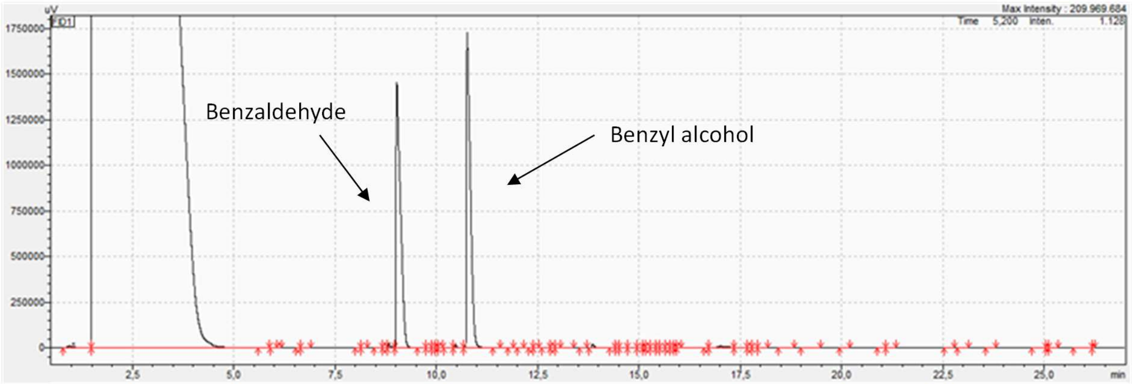


**Figure S10. GC chromatography of GtAOx substrate benzyl alcohol and product benzaldehyde.**

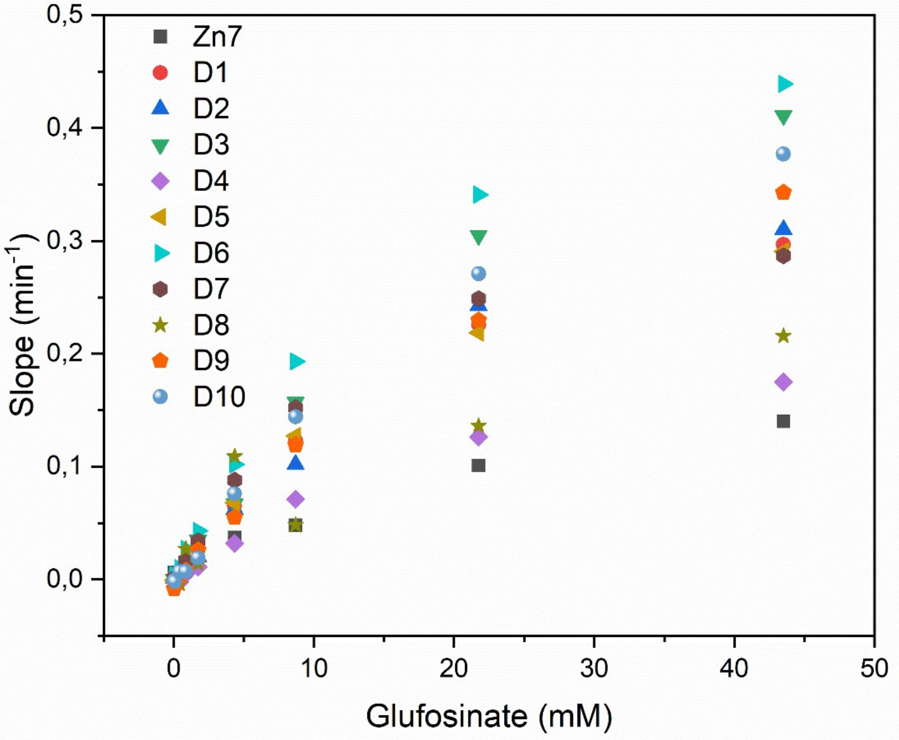


**Figure S11. Kinetics curve of purified RgDAAO variant candidates.** Ten improved RgDAAO candidates are identified after a single round of screening.

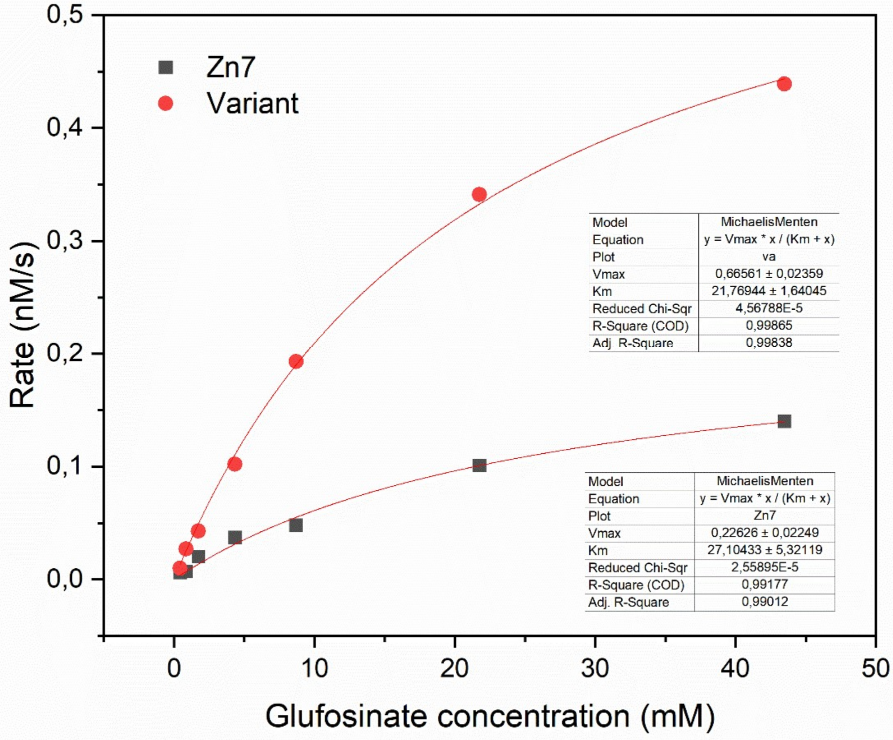


**Figure S12. Kinetics curve of RgDAAO (Parent Zn7 and variant D6) fitted with Michaelis-Menten.**

**Table S1. Primers used for library generation of GalOx, RgDAAO, and GtAOx.**

| Name | Sequence |
|---|---|
| GalOx_epPCR_F | CATATGGCAAGCGCACCGATTGGTAG |
| GalOx_epPCR_R | CTCGAGTGCGGCCGCTTGGGTAAC |
| RgDAAO_epPCR_F | CCTCTAGAAATAATTTTGTTTAACTTTAAGAAGGAGATATACCATG |
| RgDAAO_epPCR_R | GTGGTGCAGCTTACTTTCACGAG |
| GtAOx_epPCR_F | GCGCGGCAGCCATATGGTTC |
| GtAOx_epPCR_R | GTGGTGGTGGTGCTCGAGTTAG |

**Table S2. epPCR conditions for library generation of GalOx, RgDAAO, and GtAOx.**

| Reaction condition | Volume (µL) |
|---|---|
| Template (5ng/µL) | 4 |
| dNTPs (10 mM)* | 2 |
| 10xThermoPol Reaction Buffer 10 | 10 |
| Primer forward (10 µM) 2.5 | 2.5 |
| Primer reverse (10 µM) 2.5 | 2.5 |
| 10 mM $MnCl_2$ | 0.05, 0.1, 0.2 for GalOx and GtAOx |
| | 0.1, 0.2, 0.3 for RgDAAO |
| Taq Polymerase (5 U/µL) 0.5 | 0.5 |
| dd$H_2O$ | x |

*For GalOx, unbalanced dNTPs were used with 20 mM ATP, 10 mM TTP, CTP, and GTP.

**Table S3. epPCR program for library generation of GalOx, RgDAAO, and GtAOx.**

| Steps | Temperature (°C) | Time | Cycles |
|---|---|---|---|
| Initial denaturation | 95 | 30s | 1 |
| Denaturation | 95 | 30s | 25 |
| Annealing | 60 | 30s | 25 |
| Elongation | 68 | Depend on the gene length | 25 |
| Final elongation | 68 | 10min | 1 |
| Storage | 4 | ∞ | - |

**Table S4. PCR conditions for the Megawhop cloning method.**

| Reaction condition | Volume (μL) |
|---|---|
| 2xPCRBIO VeriFi™ Mix | 25 |
| Templet (135 ng) | x |
| Megaprimers (389 ng) | y |
| dd$H_2O$ | 25-x-y |

**Table S5. PCR program for the Megawhop cloning method.**

| Steps | Temperature (°C) | Time | Cycles |
|---|---|---|---|
| Warm up | 72 | ∞ | - |
| Incubation | 72 | 5min | 1 |
| Initial denaturation | 95 | 1min | 1 |
| Denaturation | 95 | 15s | 25 |
| Annealing | 68 | 1min | 25 |
| Elongation | 72 | Depend on the gene length | 25 |
| Final elongation | 72 | 10min | 1 |
| Storage | 4 | ∞ | - |

**Table S6. Protein sequence of GalOx, RgDAAO, and GtAOx.**

| Name | Sequence |
|---|---|
| GalOx_M1 | MASAPIGSAIPRNNWAVTCDSAQSGNECNKAIDGNKDTFWHTFYGANGDPKPP<br>HTYTIDMKTTQNVNGLSVLPRQDGNQNGWIGRHEVYLSSDGTNWGSPVASGSW<br>FADSTTKYSNFETRPARYVRLVAITEANGQPWTSIAEINVFQASSYTAPQPGLGRW<br>GPTIDLPIVPAAAAIEPTSGRVLMWSSYRNDAFEGSPGGITLTSSWDPSTGIVSDRT<br>VTVTKHDMFCPGISMDGNGQIVVTGGNDAKKTSLYDSSSDSWIPGPDMQVARG<br>YQSSATMSDGRVFTIGGSWSGGVFEKNGEVYSPSSKTWTSLPNAKVNPMLTADK<br>QGLYRSDNHAWLFGWKKGSVFQAGPSTAMNWYYTSGSGDVKSAGKRQSNRGV<br>APDAMCGNAVMYDAVKGKILTFGGSPDYQDSDATTNAHIITLGEPGTSPNTVFAS<br>NGLYFARTFHTSVVLPDGSTFITGGQRRGIPFEDSTPVFTPEIYVPEQDTFYKQNPNS<br>IVRAYHSISLLLPDGRVFNGGGGLCGDCTTNHFDAQIFTPNYLYDSNGNLATRPKIT<br>RTSTQSVKVGGRITISTDSSISKASLIRYGTATHTVNTDQRRIPLTLTNNGGNSYSFQ<br>VPSDSGVALPGYWMLFVMNSAGVPSVASTIRVTQAAALEHHHHHH |
| RgDAAO | MHSQKRVVVLGSGVIGLSSALILARKGYSVHILARDLPEDVSSQTFASPWAGATWT<br>PFMTLTDGPRQAKWEESTFKKWVELVPTGHAMWLKGTRRFAQNEDGLLGHWY<br>KDITPNYRPLPSSECPPGAIGVTYDTLSVHAPKYCQYLARELQKLGATFERRTVTSLE<br>QAFDGADLVVNATGLGAKSIAGIDDQAAEPIRGQTVLVKSPCKRCTTDSSDPASPA<br>YIIPRPGGEVICGGTYGVGDWDLSVNPETVQRILKHCLRLDPTISSDGTIEGIEVLRH<br>NVGLRPARRGGPRVEAERIVLPLDRTKSPLSLGRGSARAAKEKEVTLVHAYGFSQA<br>GYQQSWGAAEDVAQLVDEAFQRYHGAARESKLHHHHHH |
| GtAOx | MVHPEEVDVIVCGGGPAGCVVAGRLAYADPNLKVMLIEGGANNRDDPWVYRPG<br>IYVRNMQRDGVNDKATFYTDTMKSSHLRGRQAIVPCANILGGGSSINFQMYTRAS<br>ASDWDDFKTEGWTCQDLLPLMKRLENYQKPVNNDTHGYDGPIAISNGGQITPLA<br>QDFLRAAHSIGVPYSDDIQDLTTAHGAEIWAKYINRHTGRRSDAATAYVHSVMDV<br>QTNLYLRTNARVSRVIFEGNKAVGVAYVPSRNRAHGGAVLETIVKARKCVVLSSGT<br>LGTPQILERSGVGNGELLKKLDIKVVSDLPGVGEQYQDHYTTLSIYRVSNDSITTDDF<br>LRGVKEVQRELFQEWETSPEKARLSSNVIDAGWKLRPTEEELKEMGPEFNELWDR<br>YFKDKPDKPVMFGSIVAGAYADHTLLPPGKYMTMFQYLEYPASRGKIHIQSTNPYK<br>EPFFDSGFMNNKADFAPIRWSYKKTREVARRMDAFRGELTSHHPHFHPASAAAT<br>RDIDIKTAKEIYPDGLTVGIHMGTWHRPSEPFDASKVHEDIKYTEEDDKAIDDWVA<br>DHVETTWHSLGTCAMKPREQGGVVDKRLNVYGTEHLKCVDLSICPDNLGTNTYSS<br>ALLVGEKGADLLCEELGLKVRVPHAPVPHAPVPTGRPATQQPKH |
